## Supplementary Figures for "Stochastic Gene Expression in Auxin Signaling in the Floral Meristem of *Arabidopsis thaliana*"

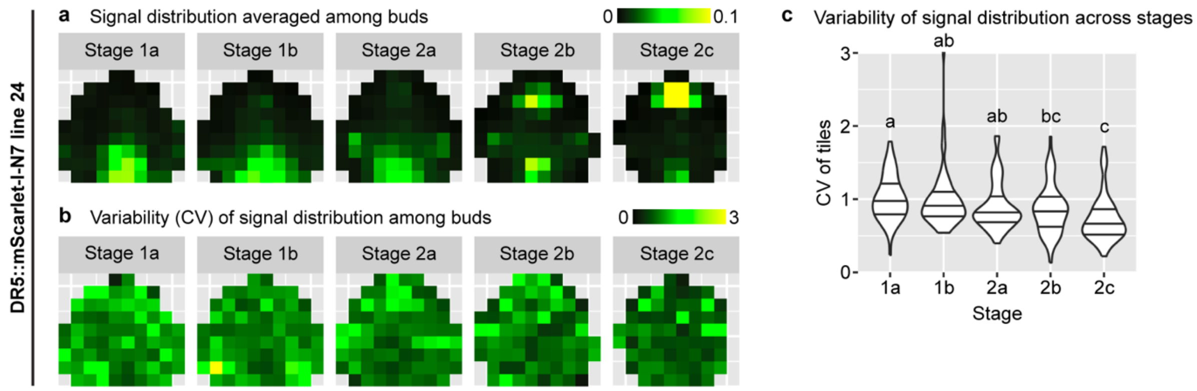


**Supplementary Fig. 1 | Analysis of an independent insertion line of *DR5::mScarlet-I-N7*.** Mean (**a**) and variability (CV) (**b-c**) of signal distribution patterns calculated from these buds: stage 1a, n = 16; stage 1b, n = 15; stage 2a, n = 21; stage 2b, n = 10; stage 2c, n = 6. In (**c**), each data point is a CV value at an (x,y) tile of a given stage. Lines show quartiles. Letters denote statistical significance in pairwise two-sided permutation tests with Bonferroni's p-value adjustments. Note that variability of global DR5 pattern decreases with developmental stage. Related to Fig. 2.

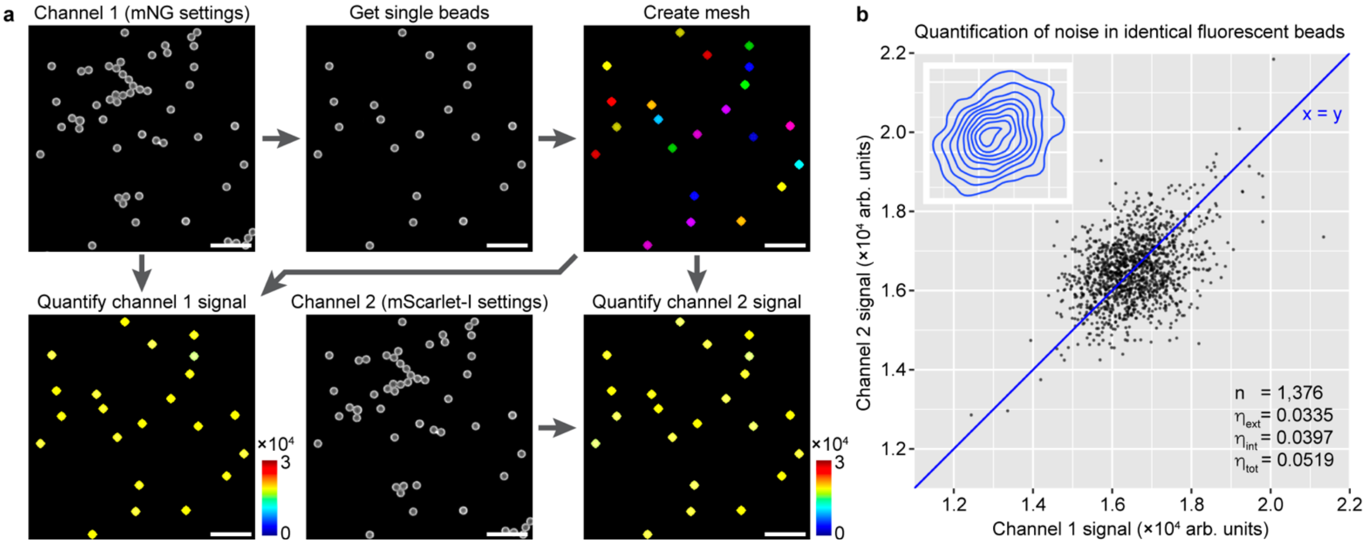


**Supplementary Fig. 2 | Quantification of extrinsic and intrinsic noise caused by instrument and measurement errors. a** Experimental procedure. Fluorescent beads were imaged using mNG and mScarlet-I settings. Single beads were used to create meshes, which were then used to quantify signal from both channels. Scale bars, 20 μm. **b** Dot plot and quantification of extrinsic ($\eta_{ext}$), intrinsic ($\eta_{int}$), and total noise ($\eta_{tot}$) from n = 1,376 beads. Inset shows the distribution of dots visualized by a 2D density plot.

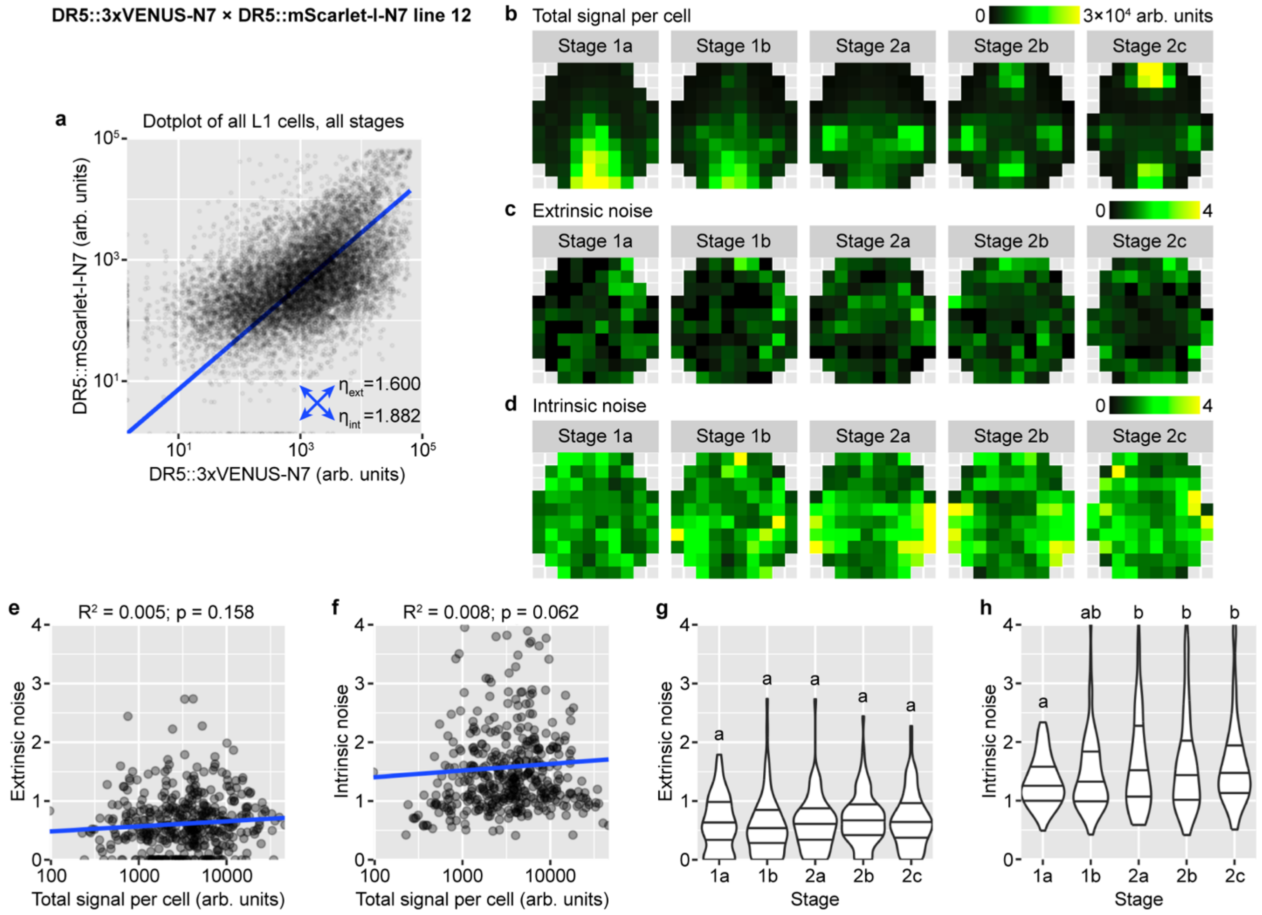


**Supplementary Fig. 3 | Analysis of an independent dual reporter line of DR5. a** Dot plot of dual-reporter signals of all cells from buds of all stages. Number of buds: stage 1a, n = 23; stage 1b, n = 16; stage 2a, n = 13; stage 2b, n = 10; stage 2c, n = 8. **b** Summed signal from both channels, averaged across all cells in each tile. **c-d** Extrinsic (**c**) and intrinsic (**d**) noise calculated from all cells in each tile, which lack apparent spatial patterns. **e-h** Relation of extrinsic and intrinsic noise to total signal per cell (**e-f**) and developmental stage (**g-h**). Each data point is a noise value at an (x,y) tile of a given stage. In (**g-h**), lines show quartiles; letters denote statistical significance in pairwise two-sided permutation tests with Bonferroni's p-value adjustments. In contrast to variability in global DR5 pattern across buds which decreases from stage 1 to 2, cellular noise either does not change (extrinsic noise) or slightly increases (intrinsic noise) with developmental stage. Related to Fig. 4.

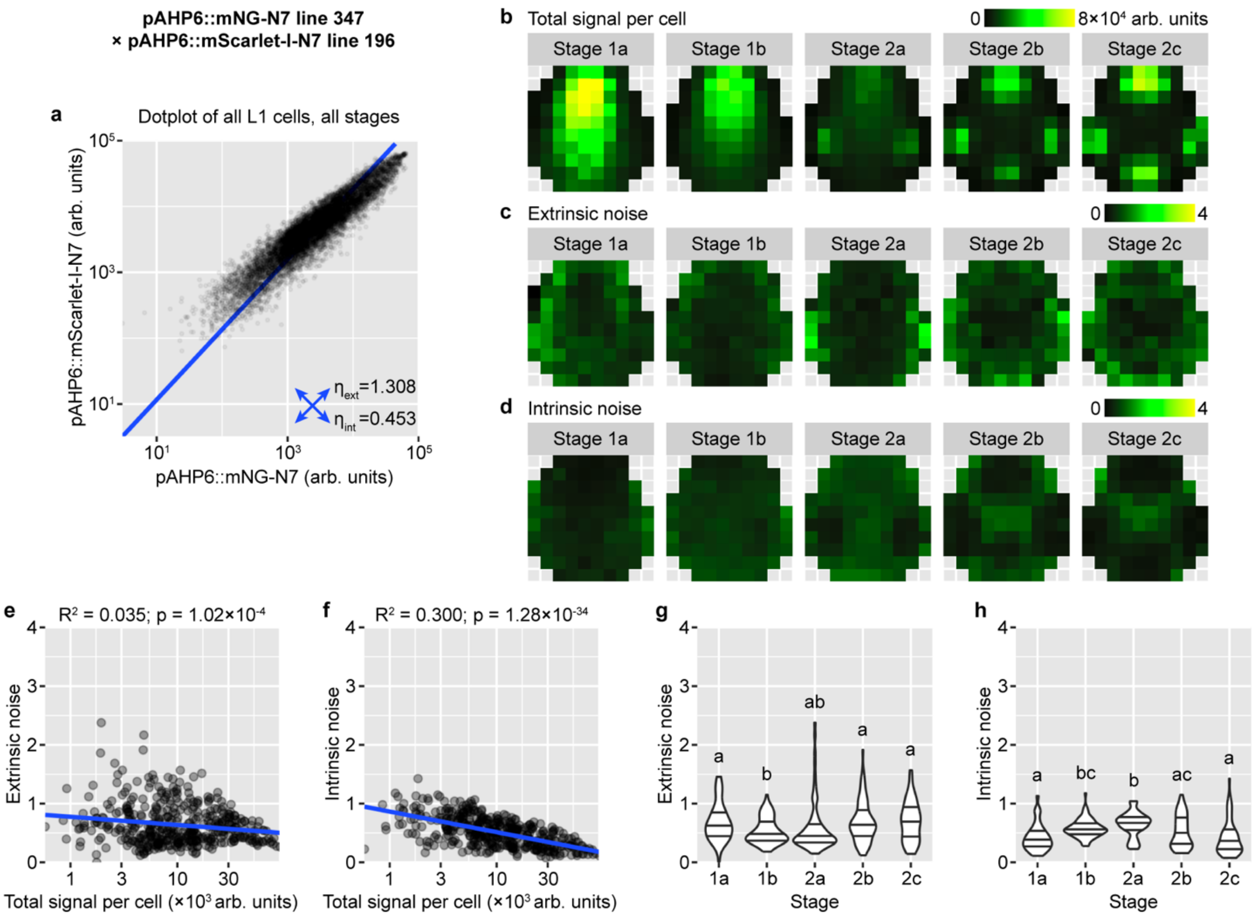


**Supplementary Fig. 4 | Analysis of an independent dual reporter line of *AHP6*. a** Dot plot of dual-reporter signals of all cells from buds of all stages. Number of buds: stage 1a, n = 14; stage 1b, n = 14; stage 2a, n = 12; stage 2b, n = 10; stage 2c, n = 6. **b** Summed signal from both channels, averaged across all cells in each tile. **c-d** Extrinsic (**c**) and intrinsic (**d**) noise calculated from all cells in each tile. Note that extrinsic noise is higher in the peripheral zone than in the central zone; intrinsic noise is lower in the incipient sepals than non-sepal regions. **e-f** Extrinsic and intrinsic noise are negatively correlated with total signal per cell. **g-h** Relation of extrinsic and intrinsic noise to developmental stage. Note that stages 1b and 2a has low extrinsic noise and high intrinsic noise than the rest of the stages. In (**e-h**), each data point is a noise value at an (x,y) tile of a given stage. In (**g-h**), lines show quartiles; letters denote statistical significance in pairwise two-sided permutation tests with Bonferroni's p-value adjustments. Related to Fig. 5.

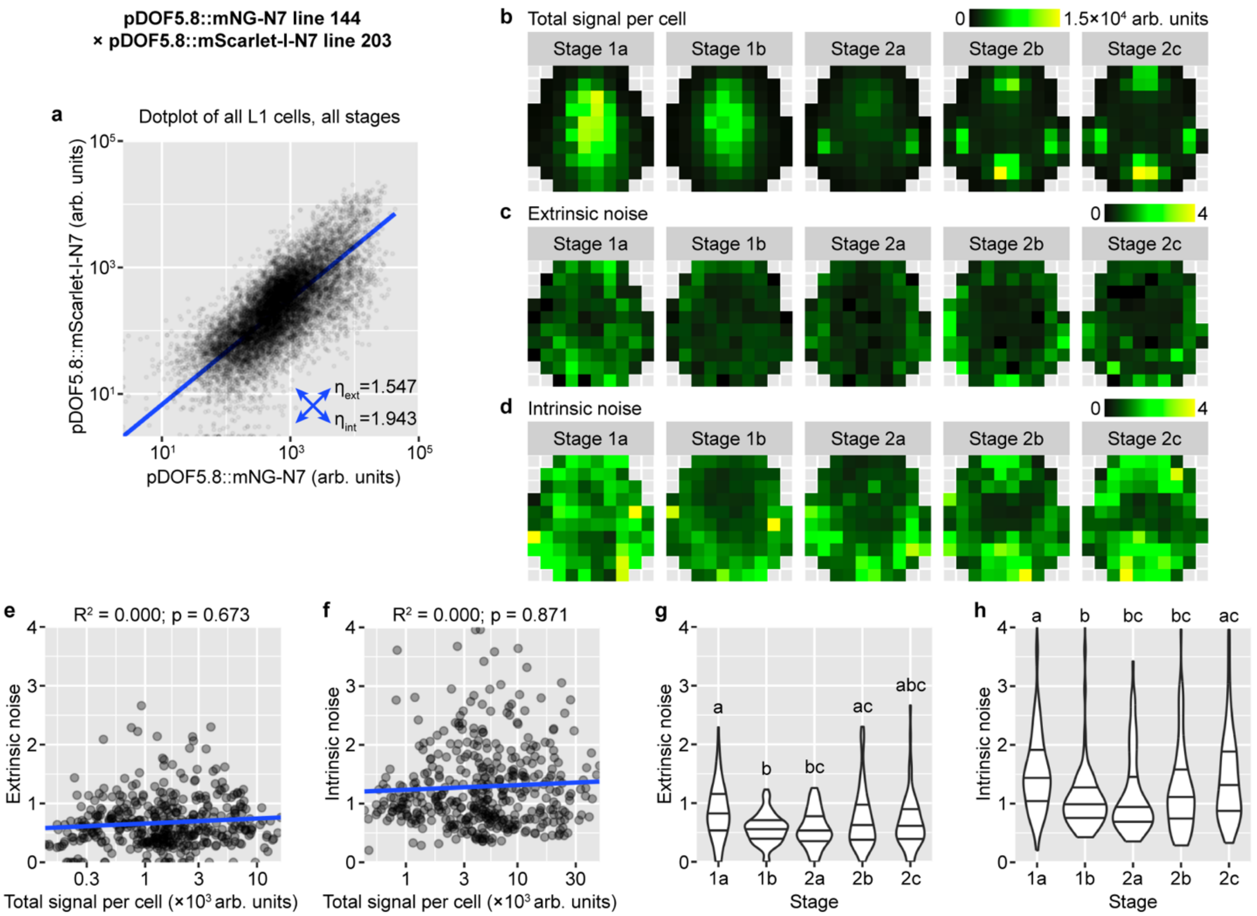


**Supplementary Fig. 5 | Analysis of an independent dual reporter line of *DOF5.8*. a** Dot plot of dual-reporter signals of all cells from buds of all stages. Number of buds: stage 1a, n = 13; stage 1b, n = 17; stage 2a, n = 10; stage 2b, n = 10; stage 2c, n = 10. **b** Summed signal from both channels, averaged across all cells in each tile. **c-d** Extrinsic (**c**) and intrinsic (**d**) noise calculated from all cells in each tile. Note that extrinsic noise is higher in the peripheral zone than in the central zone; intrinsic noise is higher in the incipient sepals than non-sepal regions. **e-f** Extrinsic and intrinsic noise are not correlated with total signal per cell. **g**-**h** Relation of extrinsic and intrinsic noise to developmental stage. Note that extrinsic and intrinsic noise decrease from stage 1a to 2a and then increase in stage 2b and 2c. In (**e-h**), each data point is a noise value at an (x,y) tile of a given stage. In (**g-h**), lines show quartiles; letters denote statistical significance in pairwise two-sided permutation tests with Bonferroni's p-value adjustments. Related to Fig. 6.

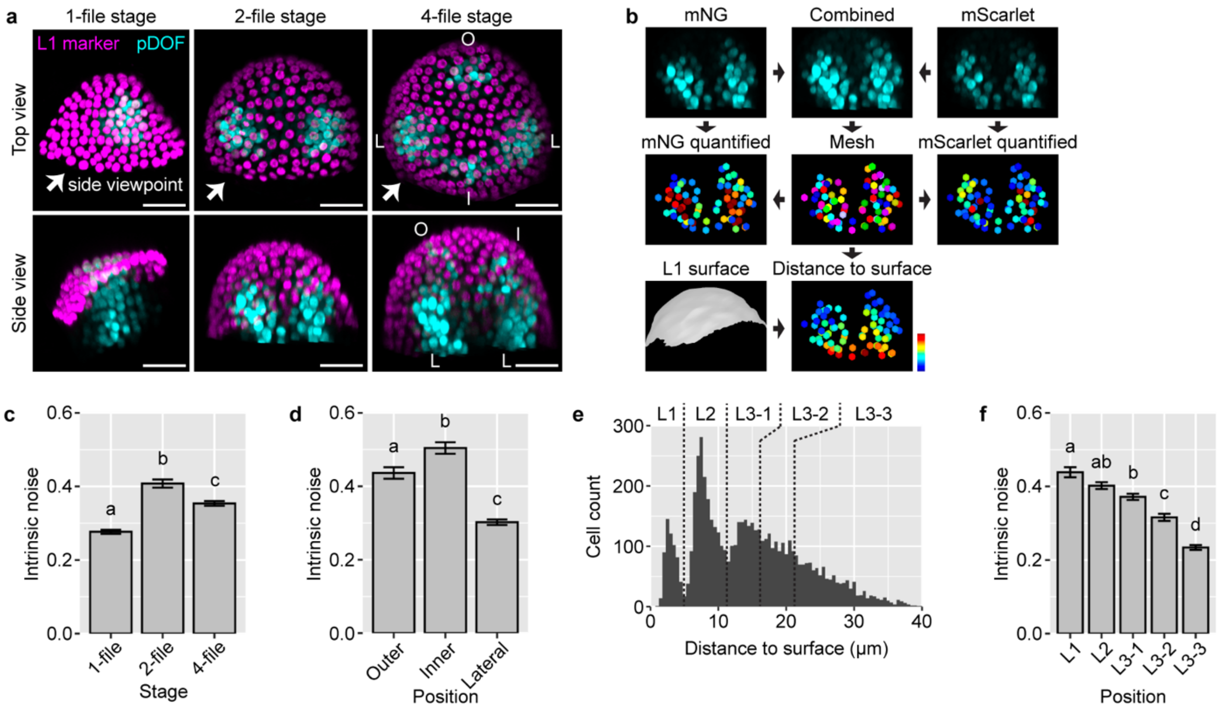


**Supplementary Fig. 6 | Analysis of the *DOF5.8* dual reporter in all cell layers reveals positional dependency of intrinsic noise. a** Representative images of *DOF5.8* expression pattern across stages. Magenta, *pATML1::H2B-TFP*. Cyan, combined signal from *pDOF5.8::mNG-N7* and *pDOF5.8::mScarlet-I-N7*. 1-file stage roughly corresponds to stage 1a to 1b; 2-file stage roughly corresponds to stage 1b to 2a; 4-file stage roughly corresponds to stage 2a to 2c. In 4-file stage, letters denote *DOF5.8*-expressing cell files in the incipient outer (O), inner (I), and lateral (L) sepal regions. Scale bars, 20 μm. **b** Workflow for quantifying the *DOF5.8* dual reporter in all cell layers. Signal from both *DOF5.8* reporters were combined, and nuclei with signal from either reporter were used to create the nuclear mesh. The nuclear mesh was used to quantify signal from either channel, with which intrinsic noise in gene expression was calculated. Distance of each nucleus to the bud surface was calculated using a surface mesh made from the epidermal marker *pATML1::H2B-TFP*. **c** Intrinsic noise is highest in the 2-file stage. **d** For buds in the 4-file stage, intrinsic noise is higher in the incipient outer and inner sepals than the incipient lateral sepals. **e** Histogram of distances from nuclei to the bud surface. Nuclei were divided into L1 (0-5 μm), L2 (5-11 μm), and artificially, L3-1 (11-16 μm), L3-2 (16-21 μm), and L3-3 (>21 μm). **f** Intrinsic noise decreases with depth in the tissue. Number of buds quantified: 1-file stage, n = 25; 2-file stage, n = 12; 4-file stage, n = 18. In (**c**, **d**, **f**), bars show intrinsic noise, and error bars show bootstrap-estimated standard deviation. Letters indicate statistical significance in two-sided permutation tests (labels of nuclei were permuted) with Bonferroni's p-value adjustment. Related to Fig. 6.

a

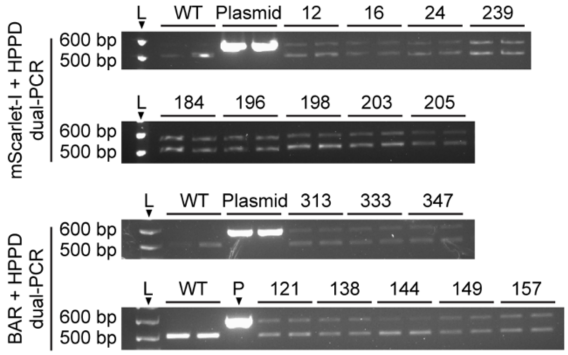


b

| Reporter | Line | Segregation ratio | Used in Figure |
|---|---|---|---|
| DR5::mScarlet-I-N7 | 239 | 194:51 (3.80:1) | 1, 2 |
| DR5::mScarlet-I-N7 | 24 | 73:27 (2.70:1) | 4, 7, S1 |
| DR5::mScarlet-I-N7 | 12 | 178:66 (2.70:1) | S2 |
| pAHP6::mNG-N7 | 313 | 103:40 (2.58:1) | 5 |
| pAHP6::mNG-N7 | 347 | 104:40 (2.60:1) | S3 |
| pAHP6::mScarlet-I-N7 | 184 | 73:27 (2.70:1) | 5 |
| pAHP6::mScarlet-I-N7 | 196 | 150:52 (2.88:1) | S3 |
| pDOF5.8::mNG-N7 | 157 | 180:63 (2.86:1) | 6, S5 |
| pDOF5.8::mNG-N7 | 144 | 75:26 (2.88:1) | S4 |
| pDOF5.8::mScarlet-I-N7 | 205 | 71:29 (2.45:1) | 6, S5 |
| pDOF5.8::mScarlet-I-N7 | 203 | 78:22 (3.55:1) | S4 |

**Supplementary Fig. 7 | Confirmation of single insertions in reporter lines. a** Dual-PCR of T1 plants. Band intensities of the transgenes (*mScarlet-I*, 609 bp, top, and *BAR*, 630 bp, bottom) were comparable to or slightly weaker than band intensities of the endogenous single-copy gene (*HPPD*, 520 bp), indicating single insertion. L, ladder. **b** Segregation ratio of antibiotic resistance (Kanamycin for mScarlet-I lines and Basta for mNG lines) in T2 seedlings. Related to Methods.
