## Supplementary figures and images for "Stochastic Gene Expression in Auxin Signaling in the Floral Meristem of *Arabidopsis thaliana*"

### 2024-05-16 dual PCR mScarlet bottom.PNG

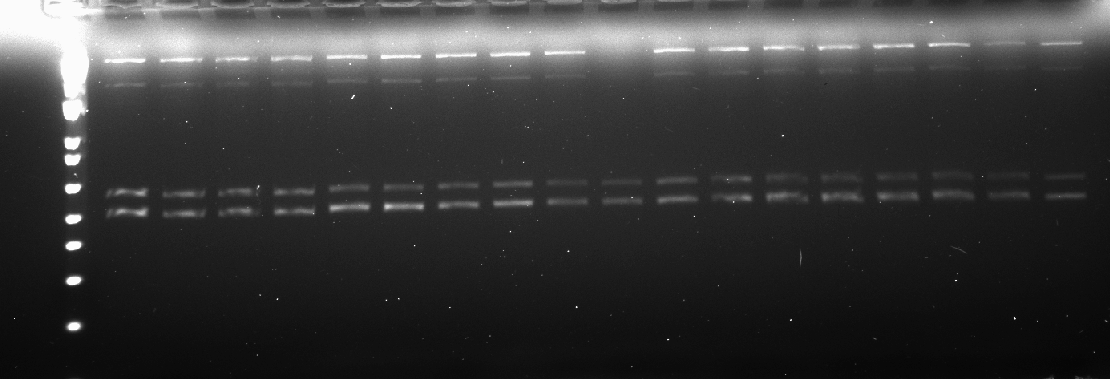

### 2024-05-16 dual PCR mScarlet top.PNG

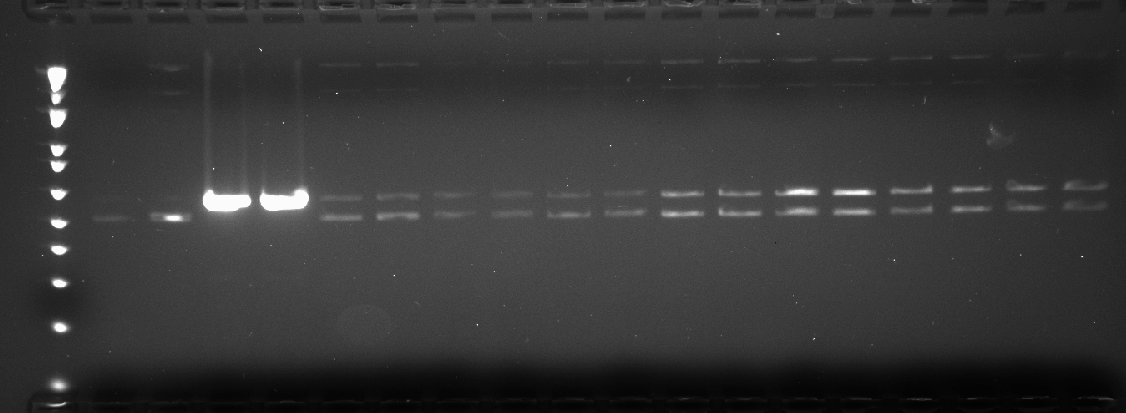

### 2024-05-16 dual PCR pAHP6-mNG.PNG

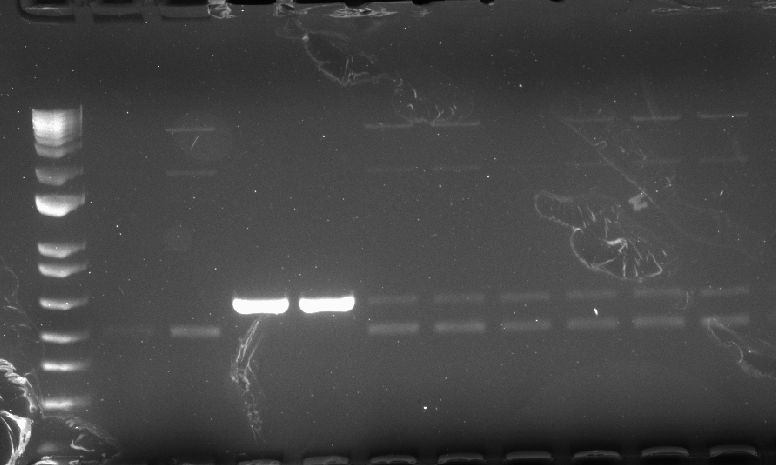

### 2024-05-16 dual PCR pDOF58-mNG.PNG

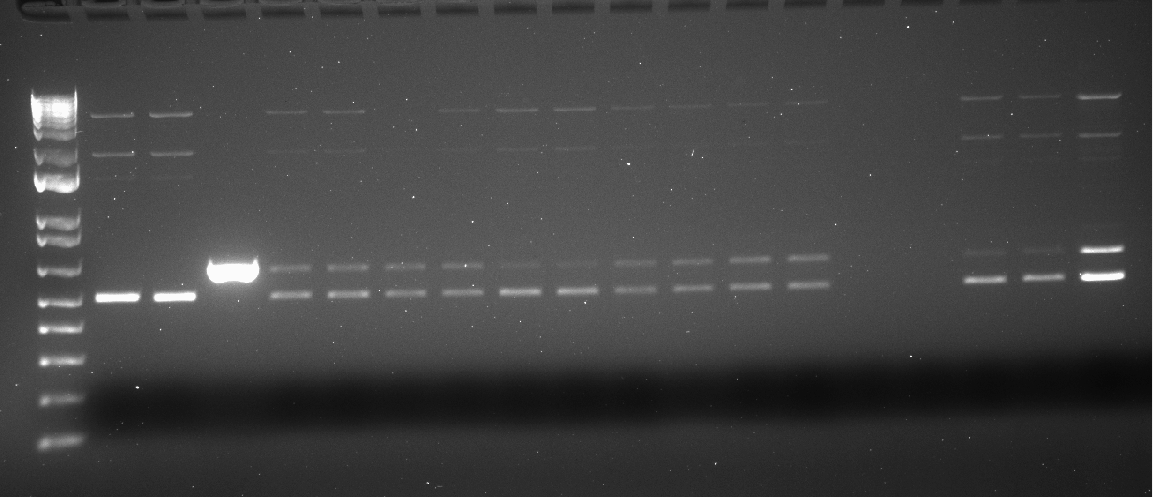
